# The Systematic Optimization of Square Wave Electroporation for Six Commonly Used Human Cell Lines

**DOI:** 10.1101/2022.05.03.490497

**Authors:** Christian Vieira, Thomas Nesmith, Saujanya Acharya, Gagan D. Gupta

## Abstract

During cellular electroporation, the formation of transient pores allow for the diffusion of innately impermeable molecules. The diversity of cell and membrane structure results in unique properties with respect to sensitivity to electric fields. The growing use of human cell lines in biomedical research and technology has led to a demand for protocols that can effectively and economically perform electroporation. We electroporated six human cell lines using a fluorescent reporter to investigate the effects of pulse electric field strength, pulse duration, and DNA concentration during electroporation. It was found that the cell lines all responded to electric field strengths within 400-950V/cm with viability decreasing with increasing voltage. It was also observed that the concentration of DNA used directly impacts transfection efficiency and cell viability as well. To better characterize square wave electroporation, we adopted a model where the pulse is described by its energy density (J/L) with respect to the sample buffer volume. It was determined that the key electrical characteristics of electroporation can be generalized with this value to provide a simplified measure of pulse intensity. The resulting analysis was consistent with other models, indicating cell type specific optimal electrical and DNA concentrations.

## 1. Introduction

Human cell lines are becoming increasingly important for analyzing a variety of cellular conditions and disease states with the increase in transfection as well as gene editing technologies. However, many of these technologies are financially or technically not viable for many research teams. Furthermore, accessibility to the same equipment, and reagents is not universal. Electroporation and electrostimulation are rising as highly flexible techniques in transfecting a variety of cell lines. Despite its proliferation amongst researchers, protocols often are not easily adapted from one apparatus to another.

Electroporation is the process of using applied voltages to increase cell membrane permeability and transport molecules (1). These applied voltages are characterized by a series of electrical conditions, which determine the force of the pulse. Pulses can be delivered primarily as two different waveforms, either exponential decay, or square wave, and can be varied in direction across multiple pulses. These are referred to as unipolar (one direction), or bipolar (two directions) pulses (2). Voltage (V) or electric field strength (V/m) provides one of the primary measures of pulse intensity. Electric field strength is determined by the voltage, and cuvette gap length. Pulse duration is another key indicator of pulse intensity. This is determined by pulse capacitance in farads (F), and sample resistance in ohm (Ω). By varying these conditions, the intensity of the pulse can be optimized for a particular cell type (3-8). When the correct electric field strength threshold is achieved, an induced charge separation is produced on either side of the cell’s plasma membrane. This causes the formation of small pores in the membrane, allowing the passage of fluid, proteins, and other molecules (9). The length of time these pores exist is variable and ranges from just a few seconds to hours (10). Unfortunately, both pore size and distribution are variable between single cells as well as across individual membranes (11). Optimal transfection is achieved by maximizing cell viability, and DNA entry. This is determined by the electric pulse conditions and the concentration of DNA. Despite these difficulties, electroporation is a robust technique for the transfection of DNA, RNA and other small soluble molecules (12).

To investigate human cell line optimization, six cell lines (HEK293T, HeLa FlpN TRex, HK-2, hTERT RPE-1 p53, MDA-MB-231 and U-2 OS) were selected for use with unipolar, square wave electroporation using varying combinations of plasmid DNA concentration, pulse voltage and duration. These provided a range of cellular origin, morphology, and function within the body. Moreover, the various cell lines also exhibit varying innate transfection efficiencies as well (13). These differences inherently require different protocols to achieve the highest efficiency. However, variation in electroporation systems limits the ability of adapting protocols based on the conditions, which can be manipulated or measured. This necessitates optimization using limited information. Energy density as a generalized value for pulse intensity incorporates the key conditions described in most protocols (**Supplemental Document 1 - Energy Density**). This can be used to bridge gaps in protocols depending on the information and electroporation system available by filling in other variables by direct measurement. The aim of this study was to demonstrate the utility of energy density as a generalized measure of pulse intensity for optimizing human cell lines between variable electroporation systems.

## 2. Materials and Methods

### 2.1 Cell culture and sample preparation

The six human cell lines were selected for electroporation: HEK293T (ATCC #CRL-3216; kind gift of B. Raught), HeLa FlpN TRex (ATCC #CCL-2 derivative line; kind gift of B. Raught; hereafter referred to as HeLa), HK-2 (ATCC #CRL-2190), hTERT RPE-1 p53 -/-(ATCC #CRL-4000 derivative line; kind gift of D. Durocher, hereafter referred to as RPE-1), MDA-MB-231(ATCC #HTB-26), and U-2 OS (ATCC #HTB-96). These were seeded onto 10cm plates and incubated at 37°C as well as 5% CO_2_ in a humidified incubator in their recommended media. Plates were grown to 90% confluency prior to electroporation. Cells were washed, trypsinized for 3mins, and resuspended in 4mL of Opti-MEM (GIBCO, 31985070). Opti-MEM was selected based on recommendations by Hyder et al. (11). From the 4mL resuspension, a predetermined volume was removed, and combined with plasmid DNA based on the concentration as well as volume of cells required for the transfections. The final sample volume per electroporation was kept at 100*µ*L. The cell counts per electroporation varied by cell line as well (**Supp. Table 1**).

### 2.2 Plasmid DNA preparation

Plasmid DNA (pEGFP-N1, 5kb (Clontech, 6085-1) encoding a cytomegalovirus (CMV) driven enhanced green fluorescent protein (GFP) was used for assessment of transfection efficiency. pEGFP-N1 was transformed in *E. coli* DH5⊡ competent cells and isolated using a midi-prep kit (Takara Bio, 740420.10), and validated by gel electrophoresis.

### 2.3 Electroporation

100*µ*L of the cell suspension sample containing the desired concentration of plasmid DNA was transferred to a 2mm (Thermo Scientific Cuvette 2mm, 5520), or 4mm gap electroporation cuvette (Bio Rad Gene Pulser Cuvette 0.4cm 851146). Square wave electroporation was performed using a Gene Pulser Xcell Eukaryotic System installed with the Capacitance Extender and Pulse Controller modules (BioRad #1652661, BioRad #1652668). Our electroporation protocols were optimized using pulse duration, pulse electric field strength, plasmid DNA concentration, and cuvette size as described for each of the experiments separately in the subsequent sections. The optimal electrical conditions for each cell line were then used to calculate the optimal energy density (**Suppl. Table 2**). These calculations assumed that resistance was approximately 120Ω after measuring the resistance of 100*µ*L of cell culture suspended in Opti-MEM (Gibco) in a 4mm cuvette using the Gene Pulser (Bio-Rad). These conditions were determined based on recommendations by Hyder et al. as well as the Gene Pulser preset conditions provided (14). Energy density was adopted to more generally characterize each pulse by combining voltage, resistance, pulse duration, and sample volume. The equations used are further described in the supplemental materials (**Energy Density Supplemental Document 1)**. Each electroporation was performed at room temperature with 5µg

DNA and cell cultures sampled at 80-90% confluency. Electroporated cells were immediately transferred to a 24 well plate with 400*µ*L of DMEM media supplemented with 10% FBS.

### 2.4 Transfection analysis determination of cell viability

To assess transfection efficiency and cell viability, samples were imaged using an EVOS FL microscope with a 10x 0.25 NA objective. 24-hrs post transfection, cells were washed, fixed, and incubated with 10µM 4′,6-diamidino-2-phenylindole (DAPI) to label the nuclei. The images obtained were analyzed and quantified using Cellprofiler (**Suppl. Fig. 15**) (15). The nuclei as well as GFP thresholds used varied per image set (**Supp. Table 2**), and were determined manually through identifying the maximum pixel intensity of false positive nuclei on a control image. The total number of DAPI labelled cells and GFP positive cells were recorded to determine the transfection efficiency (**Suppl. Figs. 3-14**). To assess cell viability, the average number of cells was calculated from experimental replicates and compared with non-electroporated controls.

## 3. Results

### 3.1 Efficiency indicates cell line dependent pulse voltage and duration

A range of electric field strengths (300-600V/cm) with a constant pulse time of 10ms were tested on the six human cell lines of different tissue origin (HEK293T - embryonic kidney;

HeLa - cervical cancer; RPE-1 - retinal pigment epithelial; U-2 OS - osteoblast; HK-2 - kidney tubule; and MDA-MB-231 - breast cancer; see Methods). Simultaneously, a range of pulse durations (5-20ms) with a constant pulse electric field strength of 530V/cm was also prepared separately. 5µg of plasmid DNA was used for each electroporation experiment. Transfection efficiency and cell viability were analyzed and plotted as a function of either electric field strength, or pulse time, to find the optimal condition for each cell type. Each cell line responded to electroporation differently, indicating varying optimal conditions (**Fig. 1 & Suppl. Figs. 1 & 2**). The HK-2, MDA-MB-231, and RPE-1 cell lines had low transfection efficiencies compared to the HeLa, HEK293T and U-2 OS lines.

**Figure 1.**
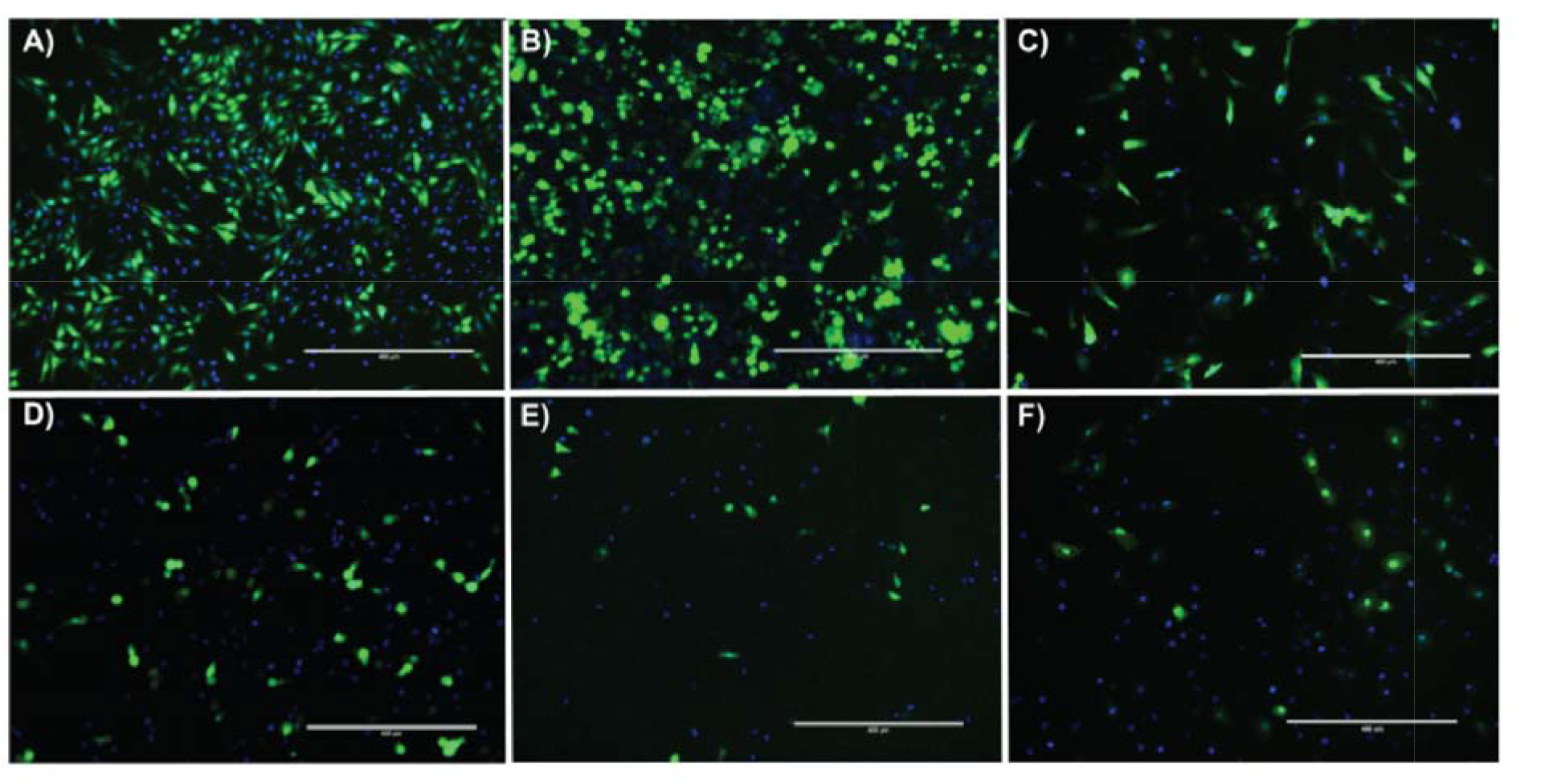
The Six Human Cell Lines Display Variable Transfection Outcomes After Electroporation is Optimized. Through 100µL electroporations using Opti-MEM and a 4mm cuvette, the effects of voltage, time and DNA Concentrations were explored. Cells were fixed and stained with 10µM DAPI(Blue). Successful electroporation was measured through GFP expression. **A)** HeLa cells electroporated at 600V/cm for 10ms with 5µg DNA. **B)** HEK 293T cells electroporated at 540V/cm for 10ms with 10µg DNA. **C)** RPE-1 cells electroporated at 600V/cm for 10ms with 10µg DNA. **D)** U-2 OS cells electroporated at 530V/cm for 10ms with 15µg DNA **E)** MDA-MB-231 cells electroporated at 600V/cm for 10ms with 15µg DNA **F)** HK-2 cells electroporated at 530V/cm for 10ms with 10µg DNA.

### 3.2 DNA concentration demonstrates logarithmic effect on transfection efficiency

DNA concentration in cell electroporation has been reported to affect transfection efficiency as well as cell survival (12). To increase transfection efficiency, DNA concentrations from 1-15µg were tested with each of the five cell lines: U-2 OS, HEK293T, HK-2, RPE-1, and MDA-MB-231. The percentage of transfected cells progressively increased as plasmid concentrations rose until it reached a peak value (**Fig. 3**). While the U-2 OS line showed peak efficiency at a lower DNA concentration of ∼3µg, the optimal concentration for the other cell lines appeared to be ∼10µg or 15µg. The HEK293T and HK-2 lines were the most responsive to increases in DNA concentration recording a ∼5-fold increase, whereas the MDA-MB-231 and U-2 OS responded poorly. As for cell viability, RPE-1 and HK-2 cell lines showed a significant improvement.

### 3.3 Optimal energy density is independent of cuvette size

It was determined that a single value for energy density can be achieved using different combinations of experimental conditions such as pulse voltage, pulse time, cuvette size and sample volume. Identical energy densities of 48kJ/L were calculated using different conditions and two different cuvette sizes: 4mm and 2mm (**Fig. 4**). The results indicated that at the lowest experimental voltage (100V), transfection efficiency dropped considerably. Conversely, cell viability was comparable to the control than higher voltage conditions. For all other conditions, the efficiency varied between 40-60% and viability also varied between 20-40%. It was also observed that optimal transfection efficiencies could be achieved regardless of cuvette type, using constant energy density (**Fig. 4**). Energy density was determined, first, by measuring the resistance of each cuvette type with a 100*µ*L buffer sample using the internal ohmmeter of the Gene Pulser (Bio-Rad). Then, the remaining electrical conditions were calculated using Equation(IX).

## 4. Discussion

### 4.1 Human cell line optimized square-wave pulse conditions vary for maximizing transfection efficiency and cell viability

Based on the electrical characteristics of a square-wave pulse, energy density was determined for each experiment using applied voltage, pulse time, capacitance, resistance, and sample volume (**Suppl. Doc. 1 Table 1**). It was observed that optimal energy density increased, or decreased proportionally with cell size (**Suppl. Doc. 1 Table 2**). This observation was supported by previous studies, which led to the development of the Schwan equation (X). This indicates the transmembrane voltage threshold for inducing permeabilization for a cell of a given size (16-19). Equation(I) (see **Supplementary Document 1**), which was used to calculate energy density, can be substituted into Equation(X) to produce Equation(XI) (see **Suppl. Fig. 16, Supplementary Document 1**). This predicts a theoretical connection between energy density and transmembrane potential, which could be used to estimate the optimal energy density conditions for future transfections.

Additionally, other factors such as cortical tension (cytoskeleton) and the phospholipid bilayer composition of cell membranes are thought to contribute towards their variable responses to the applied electric field (21,22). Transfection efficiency was observed to increase when the actin cytoskeletal network is disrupted (23). These findings suggest that cells, which readily utilize their actin networks (like highly motile cells such as MDA-MB-231), may be less receptive to electroporation, while their cytoskeletons remain intact. This agrees with our observations, where MDA-MB-231 was among the least responsive cell lines to electroporation.

It has also been found that higher concentrations of cholesterol in the membrane result in higher frequencies of pore formation (21). Conversely, anionic phosphatidylserine lipids, or membrane bound calcium ions can inhibit the formation of an electropore (24). The combined effects of these three factors are likely behind the variability we have observed between the cell lines tested. The cell lines used vary in size, cytoskeletal structure, and may vary in their membrane composition. These could all contribute to the characteristic cell-specific response to varying pulse conditions.

### 4.2 Optimal DNA concentration is cell type dependent

While our results suggest a general increase in transfection efficiency with increasing concentrations of DNA, the optimal plasmid concentration for peak transfection efficiency varied greatly between cell types. Cell lines vary in protein expression levels and tolerance to foreign DNA. For example, while the HEK293T cells in our study transfect well with higher plasmid concentrations, the U-2 OS line seems to peak at 3µg for both cell viability and transfection efficiency (**Fig. 2**). Moreover, the U-2 OS line demonstrates a significant difference in efficiency (high) and viability (low) regardless of the plasmid concentration used.

**Figure 2:**
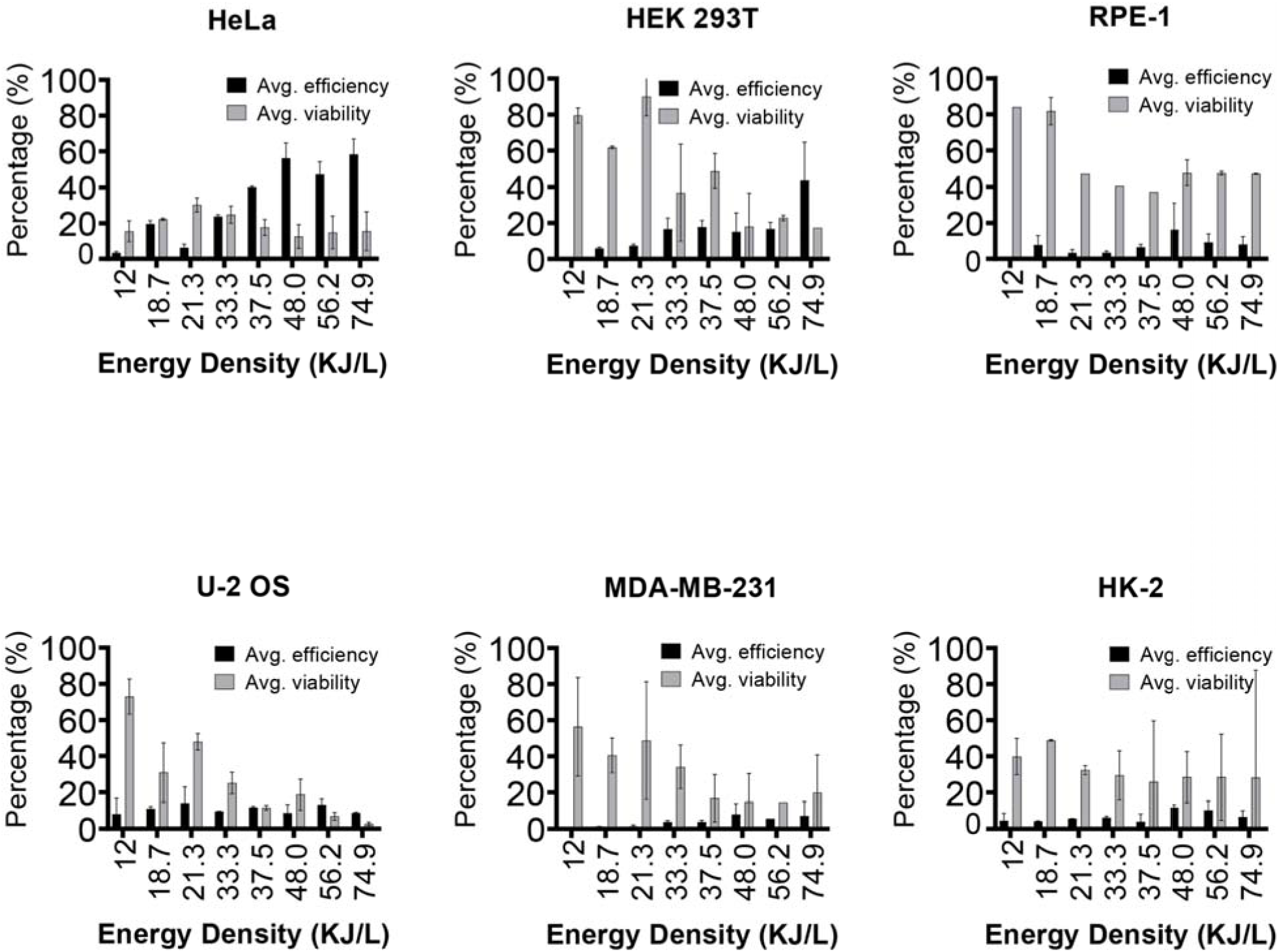
Systematic Optimization of Electroporation Parameters in Energy Density. Through 100*µ*L electroporations using Opti-MEM and a 4mm cuvette, the effects of pulse voltage and time were explored. Cells were fixed and stained with 10µM DAPI(Blue). Successful electroporation was measured through detection of GFP expression using microscopy. Microscopy images were analyzed using Cellprofiler with the conditions organized with respect to their energy densities. These Energy density values underlie the conditions as follows: 300V 10ms = 12KJ/L; 530V 5ms = 18.7KJ/L; 400V 10ms = 21.3KJ/L; 500V 10ms = 33.3KJ/L; 530V 10ms = 37.5KJ/L; 600V 10ms = 48KJ/L; 530V 15ms = 56.2KJ/L; 530V 20ms = 74.9KJ/L. HeLa, MDA-MB-231,HK-2 and HEK 293T cells responded best to electroporation conditions that applied between 48-74.9 KJ/L. RPE-1 cells respond best to 48 KJ/L electroporation conditions. U-2 OS cells responded best to 21.3 KJ/L electroporation conditions.

**Figure 3:**
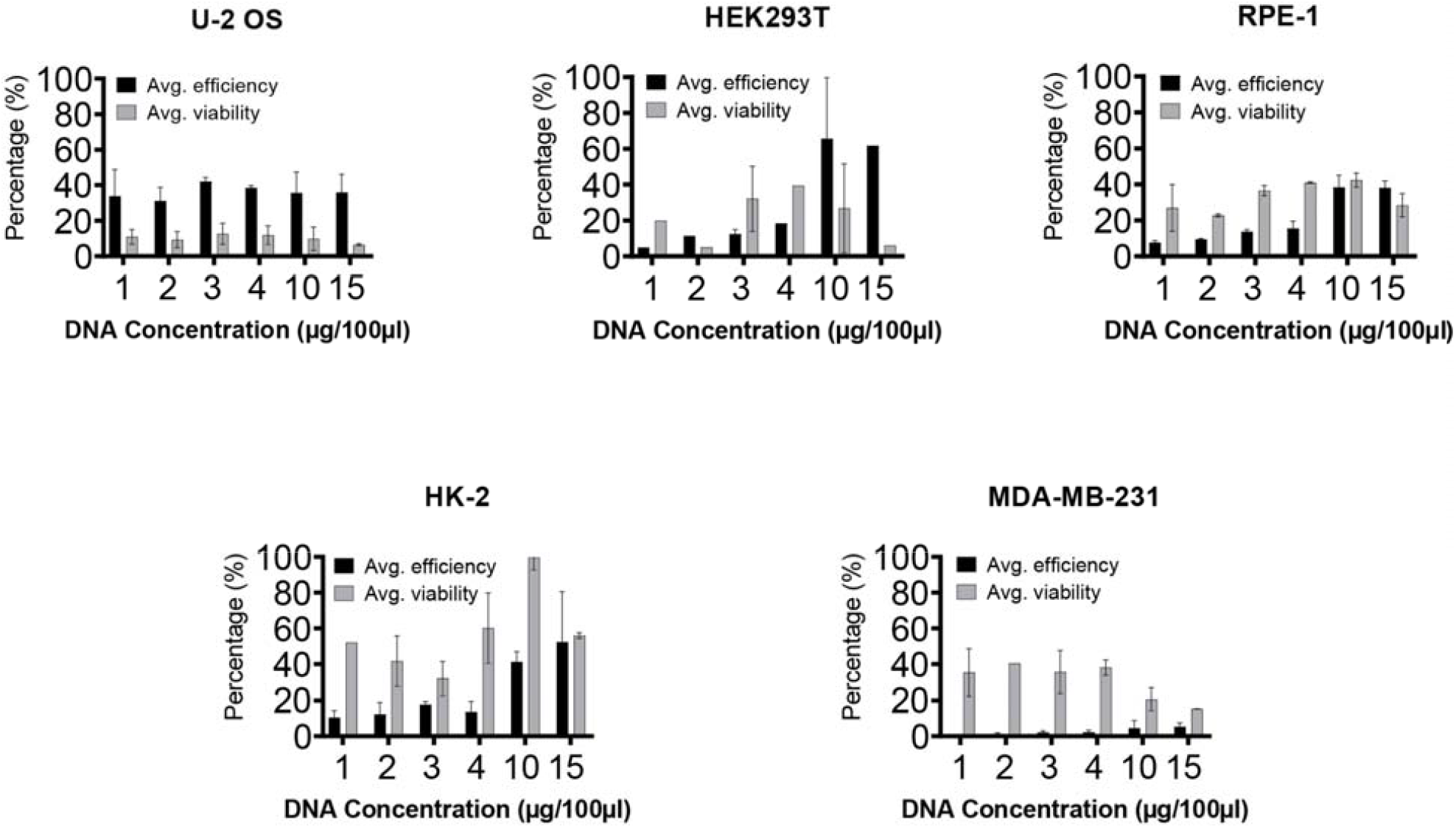
Exploring the Effect of DNA Concentration on Electroporation. Through 100µL electroporations using Opti-MEM and a 4mm cuvette, the effects of plasmid DNA concentration was examined. Cells were fixed with 4% PFA and stained with 10µM DAPI(Blue). Successful electroporation was measured through detection of GFP expression using microscopy. Microscopy images were analyzed usin Cellprofiler. Cells varied with how effectively they responded to changes in DNA concentration present during electroporation.

Additionally, these experiments were conducted with GFP, a minimally cytotoxic transfection reporter. While GFP can eventually induce apoptosis, it shows little short term cytotoxicity and transfected cells are able to survive for up to 120 hours (25). Any drop in cell viability then, would be negligible during our 48 hour experimentation period. However, overexpression of other genes can have more immediate cytotoxic effects. Therefore, determining the optimal plasmid concentration is necessary for maximizing transfection efficiency and cell viability. In general, previous reports suggest an inversely proportional relationship between increasing DNA concentration and cell viability, which is likely to occur when overexpressing a gene of interest (8,26).

### 3.3 Energy density allows for cuvette gap independent optimization

The results from **Fig. 4** indicate that when pulse duration was maximized over voltage, there was a significant drop in efficiency, but an inverse result for viability. It is likely that the transmembrane potential for the cells was not reached, based on the similar viability of the samples as compared to the control. This seems evident, given that as voltage was increased and pulse duration decreased, transfection efficiency increased proportionally. Therefore, based on this observation, maximizing voltage when calculating energy density was a more efficient way to apply the necessary threshold voltage, while achieving the same energy density. These results also indicated that for this scale of experiment, cuvette sizes were interchangeable. The 2mm cuvette provided the highest overall efficiency and viability using the optimal energy density determined experimentally with a 4mm cuvette for HEK293. However, overall the results demonstrated that approximately the same level of efficiency could be achieved with either a 2mm or 4mm respectively. This is extremely important for labs with limited resources, which may want to utilize one cuvette size to perform experiments on a variety of model cell lines. Additionally, adapting protocols from one system to another which use different shock chambers is another significant advantage provided. The use of volume allows energy density to be calculated regardless of shock chamber configuration. In turn, this enables protocols to be adapted to commercial systems or custom laboratory designs.

**Figure 4:**
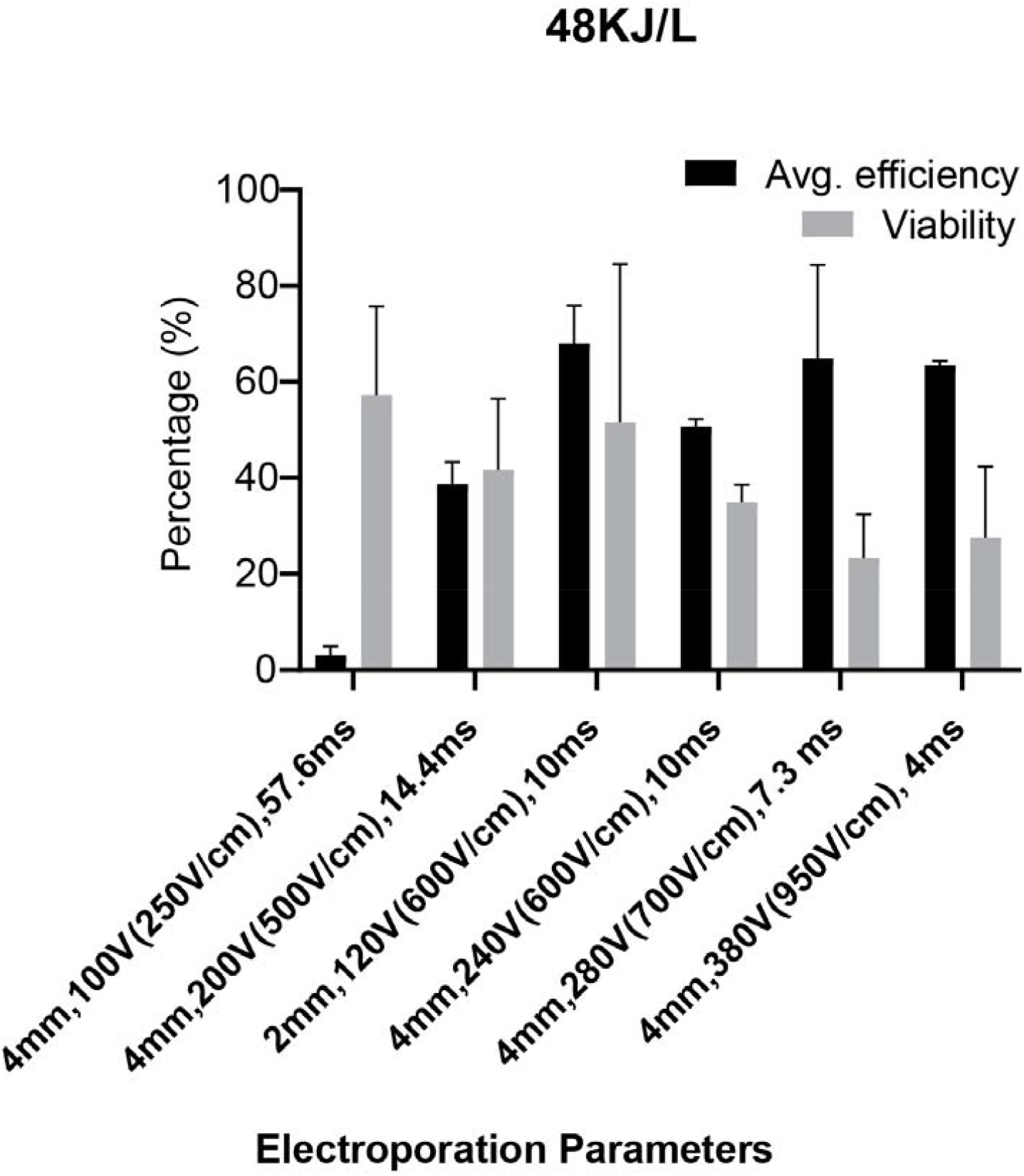
Exploration of the Effect of Pulse Duration in a Series of Energy Density Equivalent Electroporation Parameters. Through 100µL electroporations using Opti-MEM with 4-and 2-mm cuvettes, the effects of pulse duration when energy density is controlled was examined. Using an energy density constant of 48KJ/L, various parameters were derived that varied with respect to voltage and duration. Cells were fixed with 4% PFA and stained with 10µM DAPI(Blue). Successful electroporation was measured through detection of GFP expression using microscopy. Microscopy images were analyzed using Cellprofiler. As pulse duration decreased, transfection efficiency increased.

## 5. Conclusion

Our investigation demonstrated that generalizing electroporation using energy density is a highly adaptable method of transfection for human cell lines. The results support that most cell lines require specific optimization in three key areas. The first is the electrical conditions including pulse waveform, voltage and duration as well as sample buffer volume, resistance, and capacitance. These are used to calculate energy density for adapting different protocols to specific electroporation systems. The second key area is DNA concentration, which varies the level of transfection efficiency and cell viability. The third is cell type, which has the greatest impact on the optimal conditions for electroporation. For human cell lines, pulse voltage and duration for optimal transfection is closely related to cell size. Therefore, a two-fold approach is required to optimize the electroporation process for any particular experiment. The electrical conditions must be determined first. This is most reliably determined by comparing various protocols based on energy density, but could also be approximated by using cell size as an indication of the required energy density. Finally, the plasmid DNA concentration should be selected based on cell tolerance and sample volume. Efficiency still remains problematic for electroporation in many cases, and future investigations will likely reveal many of the underlying mechanisms limiting membrane permeabilization as well as trafficking of genetic material.

## Author contributions

C.V. and T.N performed all experiments, S.A., C.V. and T.N. wrote manuscript with input from G.D.G

## Acknowledgements

Profs Dan Durocher, Brian Raught and Laurence Pelletier are thanked for kind contributions of cell lines. GDG was funded by grant RGPIN-2018-04309 from NSERC.

## Tables and Legend

**Table 1:** Parameters and Results of Energy Density Equivalent Electroporation.

| Condi<br>on s | Cuvette<br>size<br>(mm) | Pulse<br>voltage<br>(Volts) | Pulse<br>time<br>(ms) | Electric<br>Field<br>Strength<br>(V/cm) | Transfection<br>Efficiency<br>(%) | Cell<br>Viabili<br>(%) |
| --- | --- | --- | --- | --- | --- | --- |
| 1 | 2 | 120 | 10 | 600 | 68.0 ±7.8 | 51.6 ±32 |
| 2 | 4 | 100 | 56 | 250 | 3.1 ±1.8 | 57.2 ±18 |
| 3 | 4 | 200 | 14.4 | 500 | 38.7 ±4.6 | 41.7 ±14 |
| 4 | 4 | 240 | 10 | 600 | 50.6 ±1.63 | 35.0 ±3 |
| 5 | 4 | 280 | 7.3 | 700 | 64.8 ±19.5 | 23.3 ±9 |
| 6 | 4 | 380 | 3 | 950 | 63.4 ±0.97 | 27.5 ±14 |

**Table 2:** Optimal Energy Densities by Cell Type.

| Energy Density (kJ/L) | Cell type (Relative Size) |
| --- | --- |
| 21.3 | U-2 OS (Large) |
| 48 | RPE-1 (Medium) |
| 48-74.9 | HeLa, MDA-MB-231, HEK 293T, HK-2 (Small) |

## References

1. Yarmush ML, Golberg A, Serša G, Kotnik T, Miklavčič D. Electroporation-based technologies for medicine: Principles, applications, and challenges. Vol. 16, Annual Review of Biomedical Engineering. 2014.

2. Orio J, Coustets M, Mauroy C, Teissie J. Electric field orientation for gene delivery using high-voltage and low-voltage pulses. J Membr Biol. 2012;245(10).

3. Knutson JC, Yee D. Electroporation: Parameters affecting transfer of DNA into mammalian cells. Anal Biochem. 1987;164(1).

4. Potter H. Electroporation in biology: Methods, applications, and instrumentation. Vol. 174, Analytical Biochemistry. 1988.

5. McCoy AM, Collins ML, Ugozzoli LA. Using the Gene Pulser MXcell electroporation system to transfect primary cells with high efficiency. J Vis Exp. 2010;(35).

6. Chicaybam L, Barcelos C, Peixoto B, Carneiro M, Limia CG, Redondo P, et al. An Efficient Electroporation Protocol for the Genetic Modification of Mammalian Cells. Front Bioeng Biotechnol. 2017;4.

7. Kotnik T, Rems L, Tarek M, Miklavcic D. Membrane Electroporation and Electropermeabilization: Mechanisms and Models. Vol. 48, Annual Review of Biophysics. 2019.

8. Hyder I, Eghbalsaied S, Kues WA. Systematic optimization of square-wave electroporation conditions for bovine primary fibroblasts. BMC Mol Cell Biol. 2020;21(1).

9. Pucihar G, Kotnik T, Miklavčič D, Teissié J. Kinetics of transmembrane transport of small molecules into electropermeabilized cells. Biophys J. 2008;

10. Teissie J. Membrane permeabilization lifetime in experiments. In: Handbook of Electroporation. 2017.

11. Golzio M, Teissié J, Rols MP. Cell synchronization effect on mammalian cell permeabilization and gene delivery by electric field. Biochim Biophys Acta - Biomembr. 2002;

12. Shi J, Ma Y, Zhu J, Chen Y, Sun Y, Yao Y, et al. A review on electroporation-based intracellular delivery. Vol. 23, Molecules. 2018.

13. Jordan ET, Collins M, Terefe J, Ugozzoli L, Rubio T. Optimizing electroporation conditions in primary and other difficult-to-transfect cells. J Biomol Tech. 2008;19(5).

14. Bio-Rad. Gene Pulser Xcell™ Electroporation System Instruction Manual. https://www.bio-rad.com/sites/default/files/webroot/web/pdf/lsr/literature/4006217A.pdf, 2021 (accessed 7 June 2021).

15. McQuin C, Goodman A, Chernyshev V, Kamentsky L, Cimini BA, Karhohs KW, et al. CellProfiler 3.0: Next-generation image processing for biology. PLoS Biol. 2018;

16. Agarwal A, Zudans I, Weber EA, Olofsson J, Orwar O, Weber SG. Effect of cell size and shape on single-cell electroporation. Anal Chem. 2007;

17. Zu Y, Huang S, Lu Y, Liu X, Wang S. Size specific transfection to mammalian cells by micropillar array electroporation. Sci Rep. 2016;

18. Marszalek P, Liu DS, Tsong TY. Schwan equation and transmembrane potential induced by alternating electric field. Biophys J. 1990;

19. Rols M-P. Parameters Affecting Cell Viability Following Electroporation In Vitro. In: Handbook of Electroporation. 2016.

21. Fernández ML, Marshall G, Sagués F, Reigada R. Structural and kinetic molecular dynamics study of electroporation in cholesterol-containing bilayers. J Phys Chem B. 2010;

22. Mescia L, Chiapperino MA, Bia P, Gielis J, Caratelli D. Modeling of electroporation induced by pulsed electric fields in irregularly shaped cells. IEEE Trans Biomed Eng. 2018;

23. Muralidharan A, Rems L, Kreutzer MT, Boukany PE. Actin networks regulate the cell membrane permeability during electroporation. Biochim Biophys Acta - Biomembr. 2021;

24. Levine ZA, Vernier PT. Calcium and phosphatidylserine inhibit lipid electropore formation and reduce pore lifetime. J Membr Biol. 2012;

25. Ansari AM, Ahmed AK, Matsangos AE, Lay F, Born LJ, Marti G, et al. Cellular GFP Toxicity and Immunogenicity: Potential Confounders in Vivo Cell Tracking Experiments. Stem Cell Reviews and Reports. 2016.

26. Ohyama H, McBride J, Wong DTW. Optimized conditions for gene transfection into the human eosinophilic cell line EoL-1 by electroporation. J Immunol Methods. 1998;

