## Supplementary Figures 1-16 for "The Systematic Optimization of Square Wave Electroporation for Six Commonly Used Human Cell Lines"

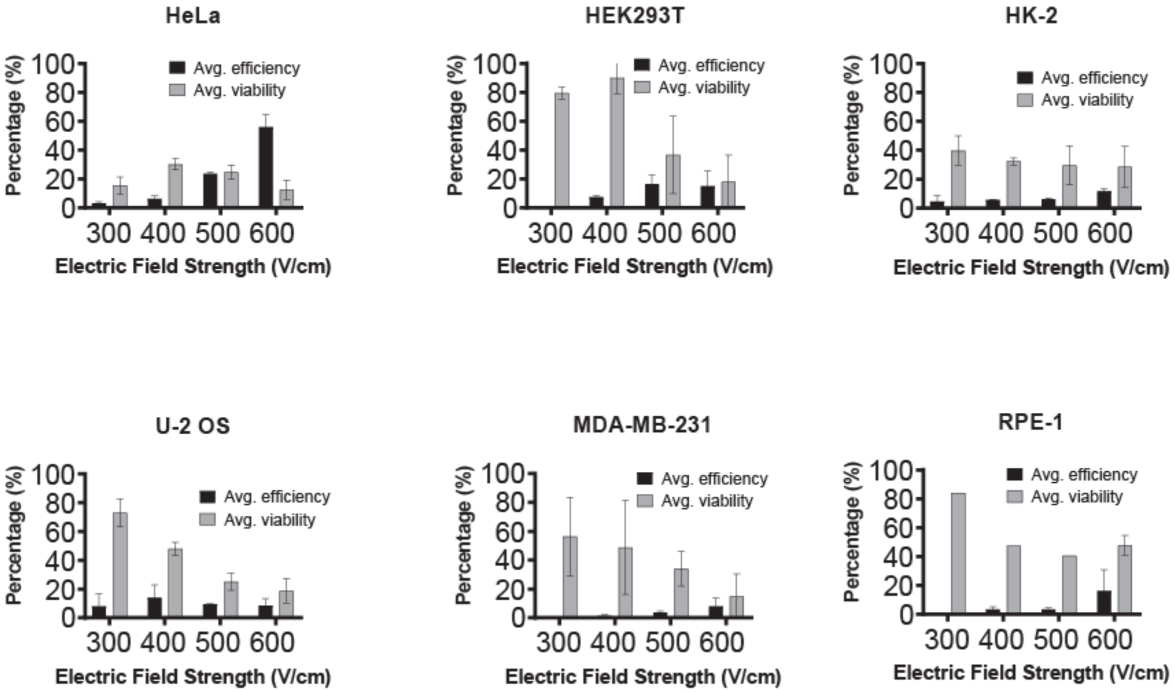


**Supplementary Figure 1: Systematic Optimization of Pulse Electric Field Strength on Electroporation.**

Through 100µl electroporations using Opti-MEM and a 4mm cuvette, the effects of pulse Electric field strengths between 300-600 V/cm was explored. Cells were fixed and stained with 10µM DAPI (Blue). Successful electroporation was measured through detection of GFP expression using microscopy. Microscopy images were analyzed using Cellprofiler.

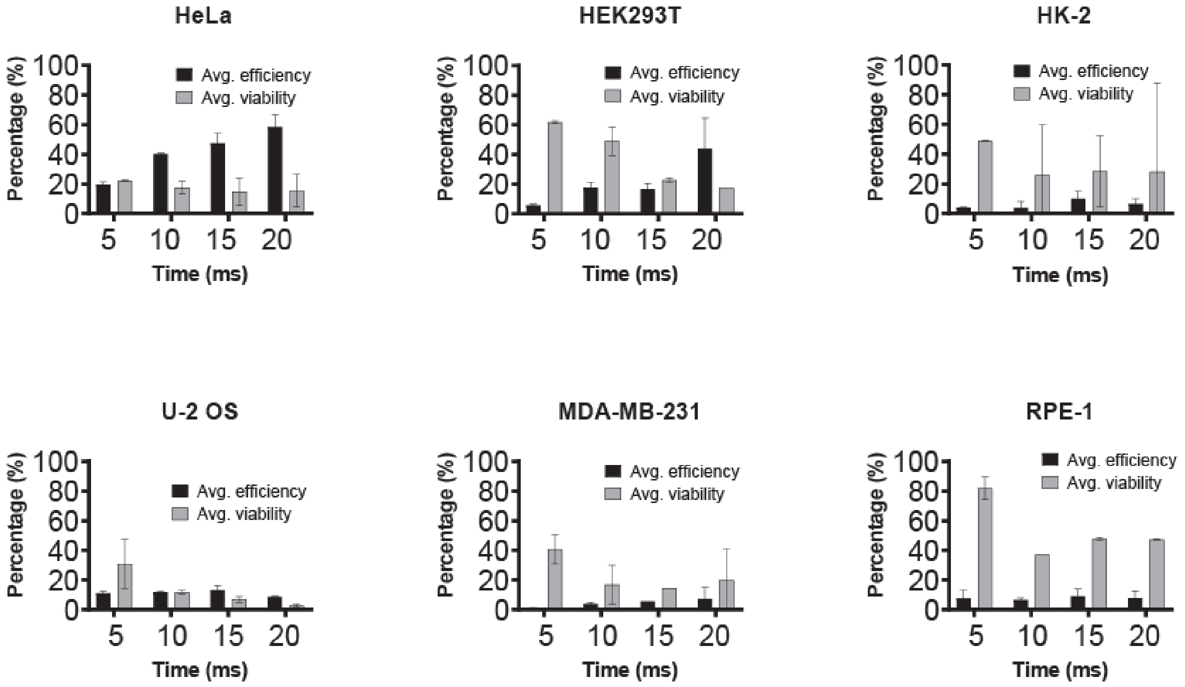


**Supplementary Figure 2: Systematic Optimization of Pulse Duration of Electroporation with Constant Electric Field Strength.**

Through 100µl electroporations using Opti-MEM and a 4mm cuvette, the effects of pulse durations between 5-20ms were explored. Cells were fixed and stained with 10µM DAPI(Blue). Successful electroporation was measured through detection of GFP expression using microscopy. Microscopy images were analyzed using Cellprofiler.

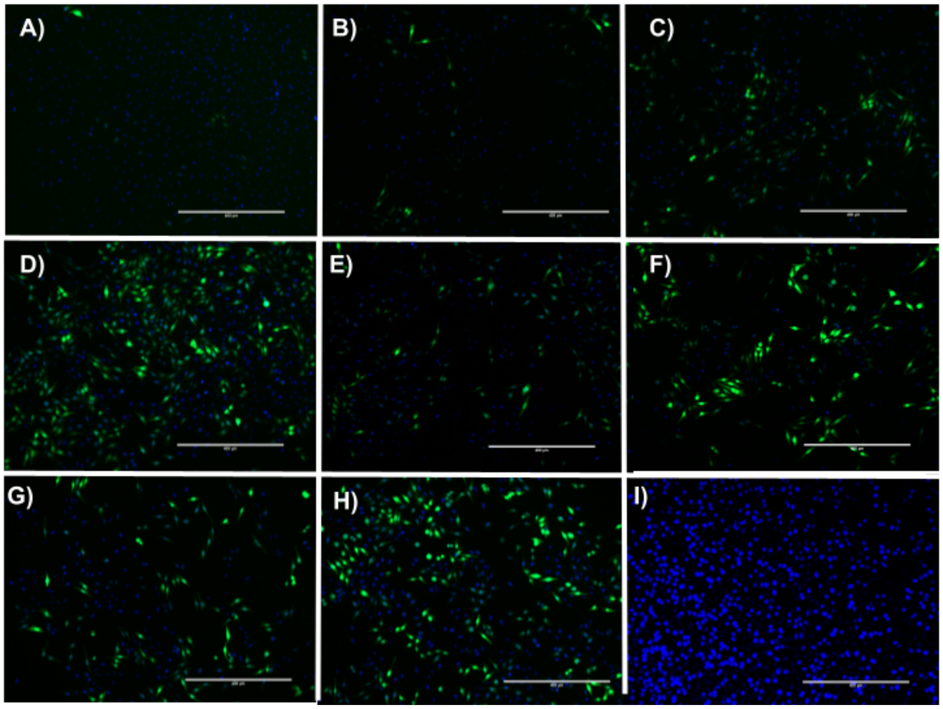


**Supplementary Figure 3. HeLa Energy Density Gradient.**

Through 100µl electroporations using Opti-MEM and a 4mm cuvette, the effects of pulse voltage and time were explored. Cells were fixed with 4% PFA and stained with 10µM DAPI(Blue). Successful electroporation was measured through detection of GFP expression using microscopy. Microscopy images were analyzed using Cellprofiler. The Experimental conditions were as follows: **A)** 4mm 300V/cm 10ms **B)** 2mm 400V/cm 10ms **C)** 4mm 500V/cm 10ms **D)** 4mm 600V/cm 10ms **E)** 4mm 530V/cm 5ms **F)** 4mm 530V/cm 10ms **G)** 4mm 530V/cm 15ms **H)** 4mm 530V/cm 20ms **I)** Control. Scale = 400µm.

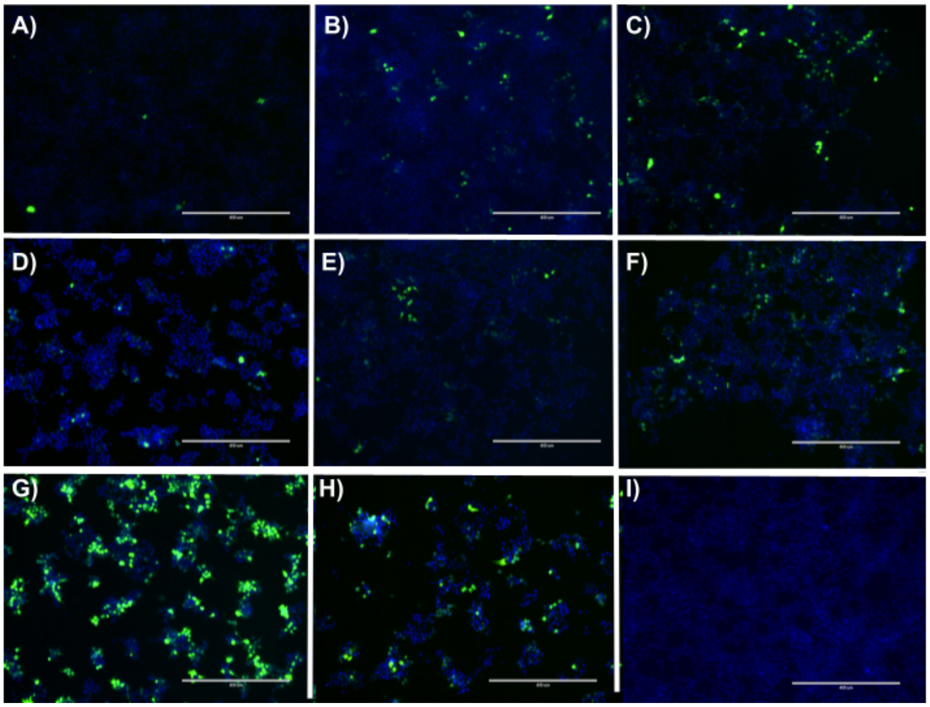


**Supplementary Figure 4. HEK Energy Density Gradient**

Through 100µl electroporations using Opti-MEM and a 4mm cuvette, the effects of pulse voltage and time were explored. Cells were fixed with 4% PFA and stained with 10µM DAPI (Blue). Successful electroporation was measured through detection of GFP expression using microscopy. Microscopy images were analyzed using Cellprofiler. The Experimental conditions were as follows: **A)** 4mm 300V/cm 10ms **B)** 2mm 400V/cm 10ms **C)** 4mm 500V/cm 10ms **D)** 4mm 600V/cm 10ms **E)** 4mm 530V/cm 5ms **F)** 4mm 530V/cm 10ms **G)** 4mm 530V/cm 15ms **H)** 4mm 530V/cm 20ms **I)** Control. Scale = 400µm.

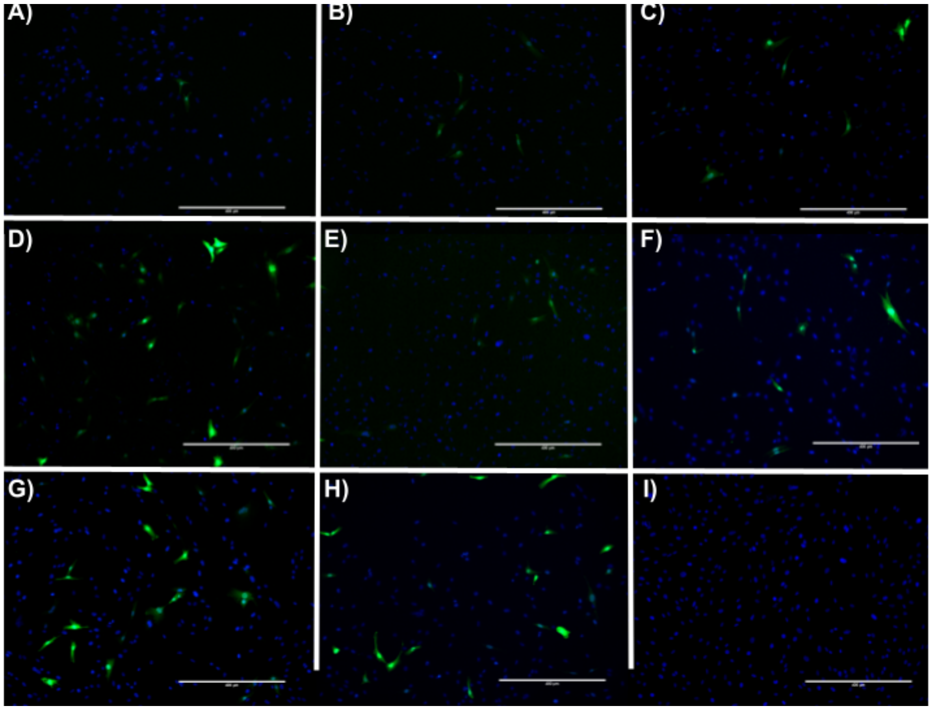


**Supplemental Figure 5. RPE-1 Energy Density Gradient.**

Through 100µl electroporations using Opti-MEM and a 4mm cuvette, the effects of pulse voltage and time were explored. Cells were fixed with 4% PFA and stained with 10µM DAPI (Blue). Successful electroporation was measured through detection of GFP expression using microscopy. Microscopy images were analyzed using Cellprofiler. The Experimental conditions were as follows: **A)** 4mm 300V/cm 10ms **B)** 2mm 400V/cm 10ms **C)** 4mm 500V/cm 10ms **D)** 4mm 600V/cm 10ms **E)** 4mm 530V/cm 5ms **F)** 4mm 530V/cm 10ms **G)** 4mm 530V/cm 15ms **H)** 4mm 530V/cm 20ms **I)** Control. Scale = 400µm.

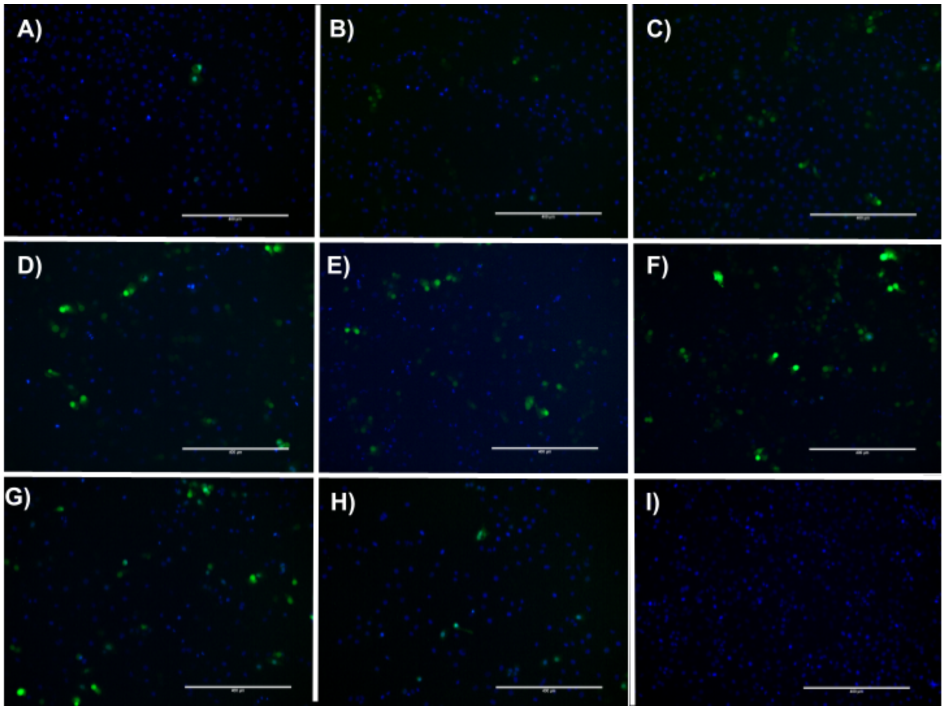


**Supplementary Figure 6. HK-2 Energy Density Gradient.**

Through 100µl electroporations using Opti-MEM and a 4mm cuvette, the effects of pulse voltage and time were explored. Cells were fixed with 4% PFA and stained with 10µM DAPI (Blue). Successful electroporation was measured through detection of GFP expression using microscopy. Microscopy images were analyzed using Cellprofiler. The Experimental conditions were as follows: **A)** 4mm 300V/cm 10ms **B)** 2mm 400V/cm 10ms **C)** 4mm 500V/cm 10ms **D)** 4mm 600V/cm 10ms **E)** 4mm 530V/cm 5ms **F)** 4mm 530V/cm 10ms **G)** 4mm 530V/cm 15ms **H)** 4mm 530V/cm 20ms **I)** Control. Scale = 400µm.

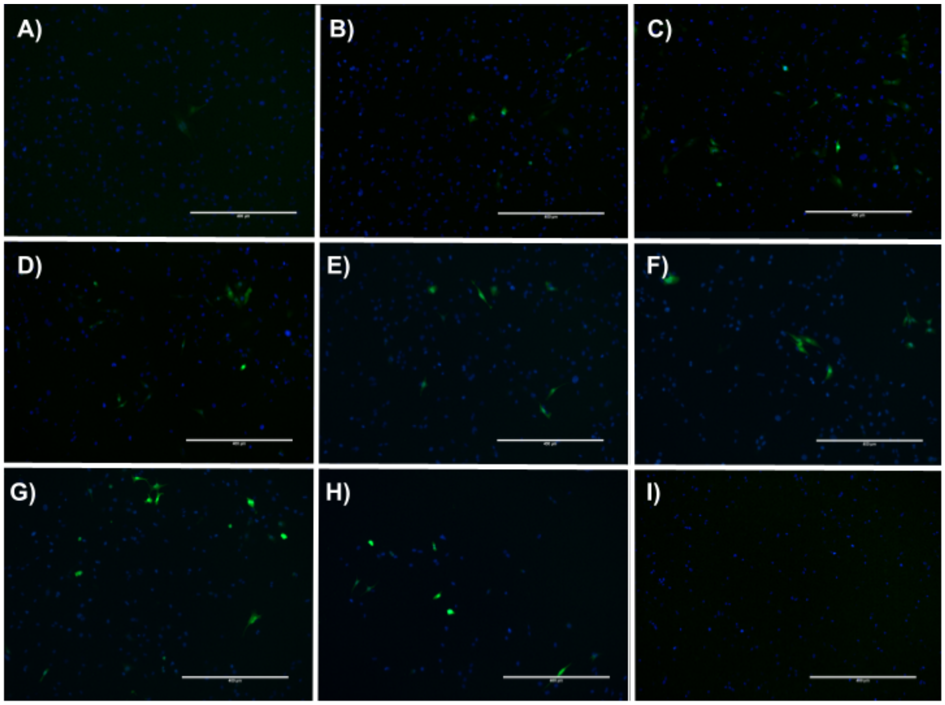


**Supplementary Figure 7. MDA-MB Energy Density Gradient.**

Through 100µl electroporations using Opti-MEM and a 4mm cuvette, the effects of pulse voltage and time were explored. Cells were fixed with 4% PFA and stained with 10µM DAPI (Blue). Successful electroporation was measured through detection of GFP expression using microscopy. Microscopy images were analyzed using Cellprofiler. The Experimental conditions were as follows: **A)** 4mm 300V/cm 10ms **B)** 2mm 400V/cm 10ms **C)** 4mm 500V/cm 10ms **D)** 4mm 600V/cm 10ms **E)** 4mm 530V/cm 5ms **F)** 4mm 530V/cm 10ms **G)** 4mm 530V/cm 15ms **H)** 4mm 530V/cm 20ms **I)** Control. Scale = 400µm.

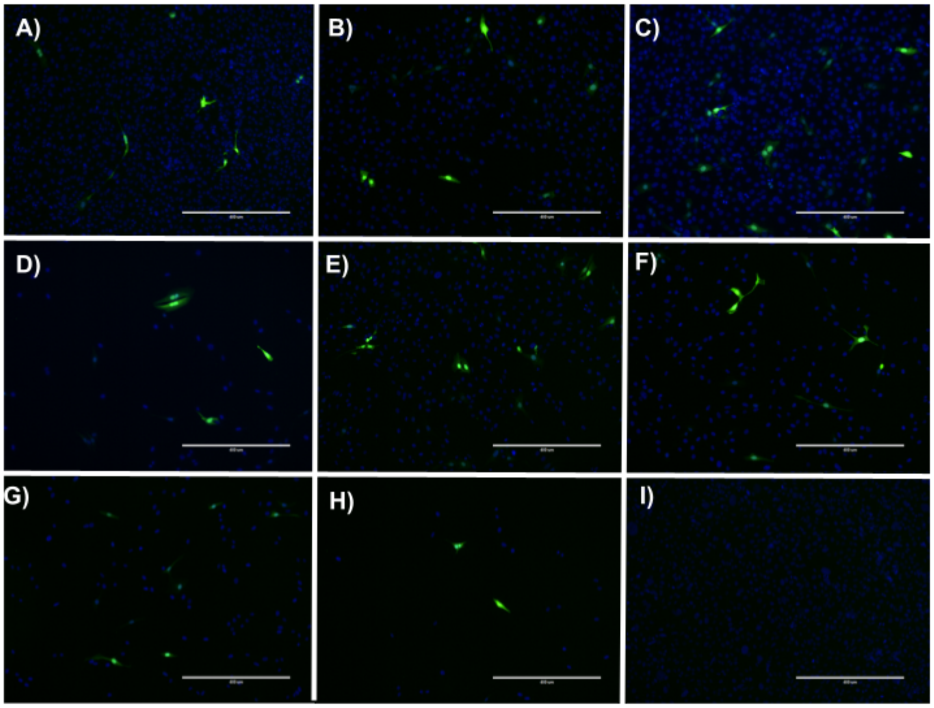


**Supplementary Figure 8. U-2 OS Energy Density Gradient.**

Through 100µl electroporations using Opti-MEM and a 4mm cuvette, the effects of pulse voltage and time were explored. Cells were fixed with 4% PFA and stained with 10µM DAPI (Blue). Successful electroporation was measured through detection of GFP expression using microscopy. Microscopy images were analyzed using Cellprofiler. The Experimental conditions were as follows: **A)** 4mm 300V/cm 10ms **B)** 2mm 400V/cm 10ms **C)** 4mm 500V/cm 10ms **D)** 4mm 600V/cm 10ms **E)** 4mm 530V/cm 5ms **F)** 4mm 530V/cm 10ms **G)** 4mm 530V/cm 15ms **H)** 4mm 530V/cm 20ms **I)** Control. Scale = 400µm.

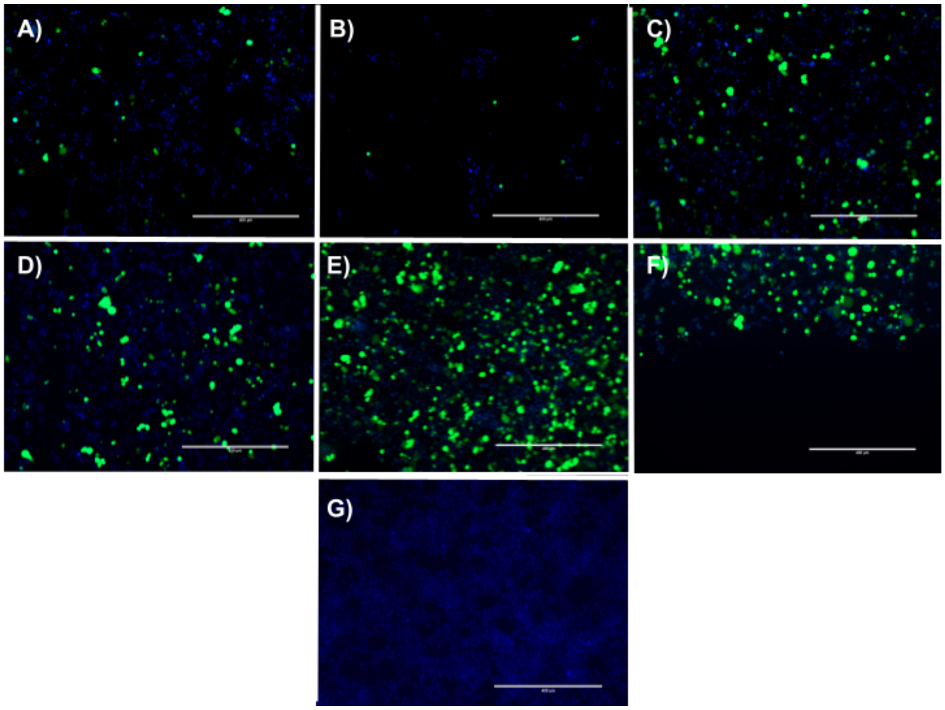


**Supplementary Figure 9. HEK293T DNA Gradient.**

Through 100µl electroporations using Opti-MEM and a 4mm cuvette, the effect of DNA concentration was explored. A constant pulse of 540V/cm for 10ms was applied to all conditions. Cells were fixed and stained with 10µM DAPI (Blue). Successful electroporation was measured through detection of GFP expression using microscopy. Microscopy images were analyzed using Cellprofiler. The Experimental conditions were as follows: **A)** 1µg **B)** 2µg **C)** 3µg **D)** 4µg **E)** 10µg **F)** 15µg **G)** Control. Scale = 400µm.

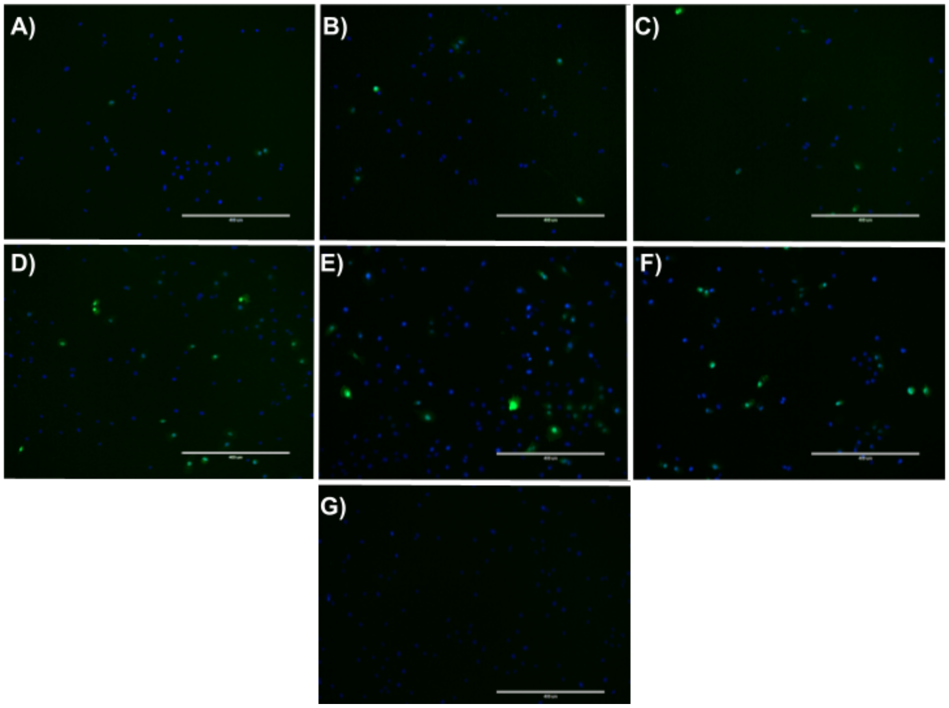


**Supplementary Figure 10. HK-2 DNA Gradient.**

Through 100µl electroporations using Opti-MEM and a 4mm cuvette, the effect of DNA concentration was explored. A constant pulse of 600V/cm for 10ms was applied to all conditions. Cells were fixed and stained with 10µM DAPI (Blue). Successful electroporation was measured through detection of GFP expression using microscopy. Microscopy images were analyzed using Cellprofiler. The Experimental conditions were as follows: **A)** 1µg **B)** 2µg **C)** 3µg **D)** 4µg **E)** 10µg **F)** 15µg **G)** Control. Scale = 400µm.

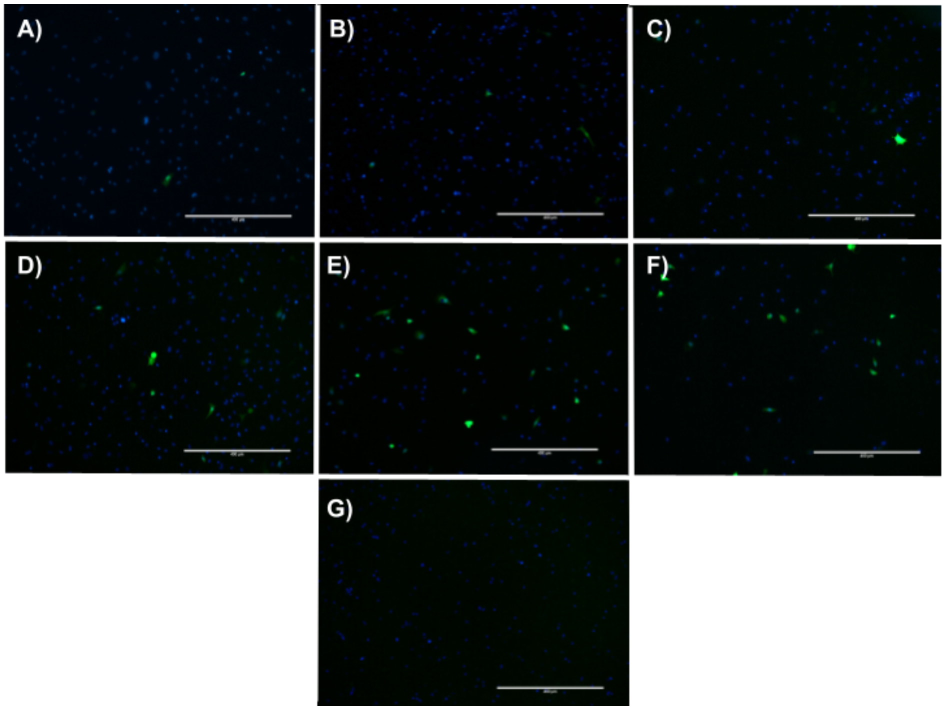


**Supplementary Figure 11. MDA-MB DNA Gradient.**

Through 100µl electroporations using Opti-MEM and a 4mm cuvette, the effect of DNA concentration was explored. A constant pulse of 600V/cm for 10ms was applied to all conditions. Cells were fixed and stained with 10µM DAPI (Blue). Successful electroporation was measured through detection of GFP expression using microscopy. Microscopy images were analyzed using Cellprofiler. The Experimental conditions were as follows: **A)** 1µg **B)** 2µg **C)** 3µg **D)** 4µg **E)** 10µg **F)** 15µg **G)** Control. Scale = 400µm.

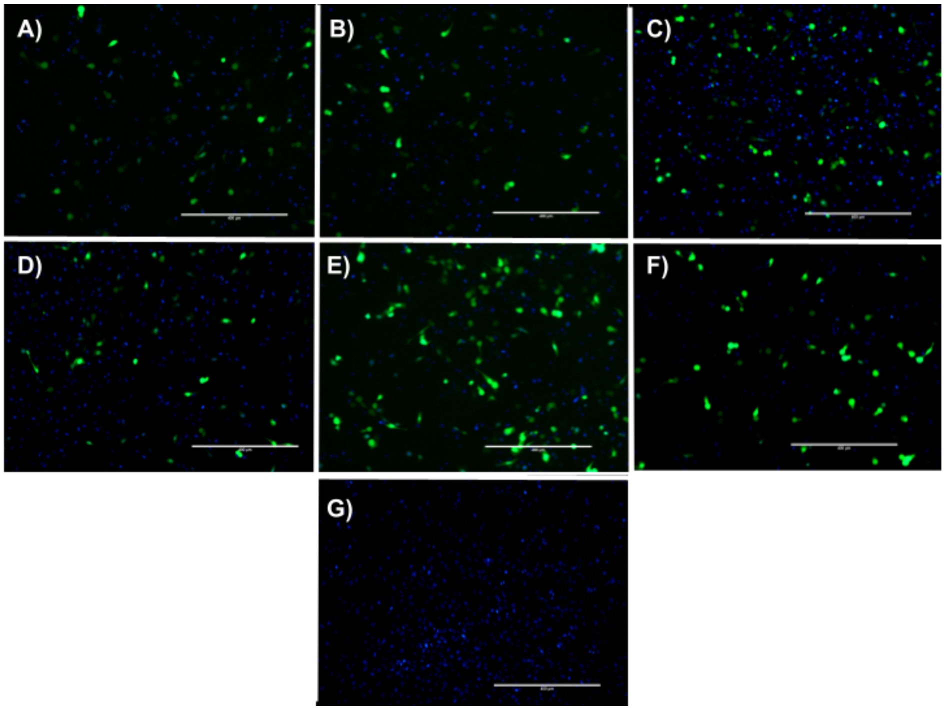


**Supplementary Figure 12. U2OS DNA Gradient.**

Through 100 µl electroporations using Opti-MEM and a 4 mm cuvette, the effect of DNA concentration was explored. A constant pulse of 530 V/cm for 10 ms was applied to all conditions. Cells were fixed and stained with 10 mM DAPI(Blue). Successful electroporation was measured through detection of GFP expression using microscopy. Microscopy images were analyzed using Cellprofiler. The Experimental conditions were as follows: **A)** 1 µg **B)** 2 µg **C)** 3 µg **D)** 4 µg **E)** 10 µg **F)** 15 µg **G)** Control. Scale = 400 µm.

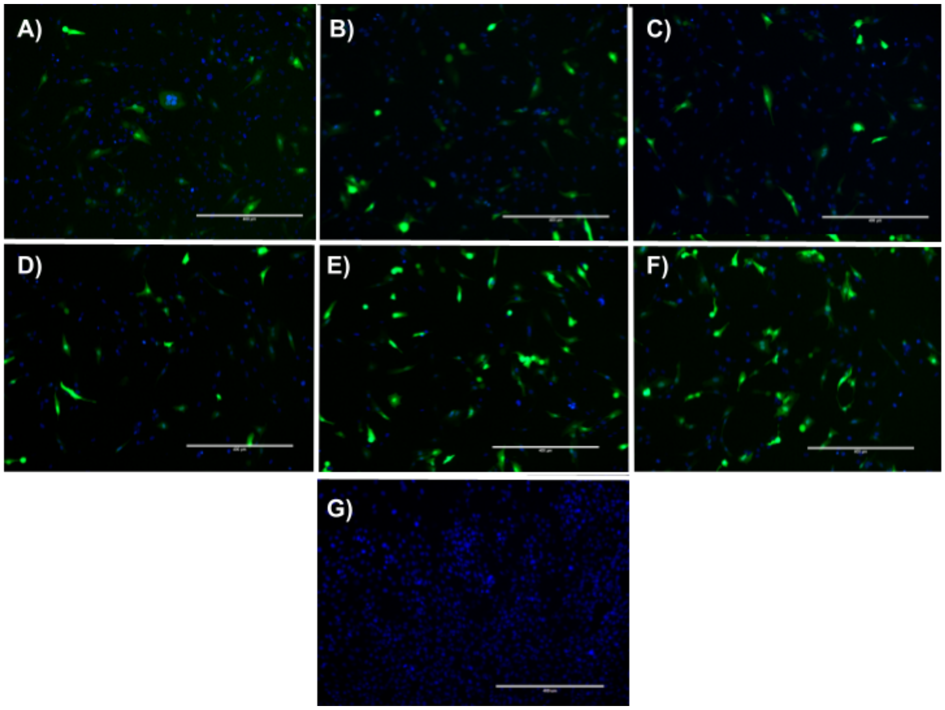


**Supplementary Figure 13. RPE-1 DNA Gradient.**

Through 100µl electroporations using Opti-MEM and a 4mm cuvette, the effect of DNA concentration was explored. A constant pulse of 600V/cm for 10ms was applied to all conditions. Cells were fixed and stained with 10µM DAPI(Blue). Successful electroporation was measured through detection of GFP expression using microscopy. Microscopy images were analyzed using Cellprofiler. The Experimental conditions were as follows: **A)** 1µg **B)** 2µg **C)** 3µg **D)** 4µg **E)** 10µg **F)** 15µg **G)** Control. Scale = 400µm.

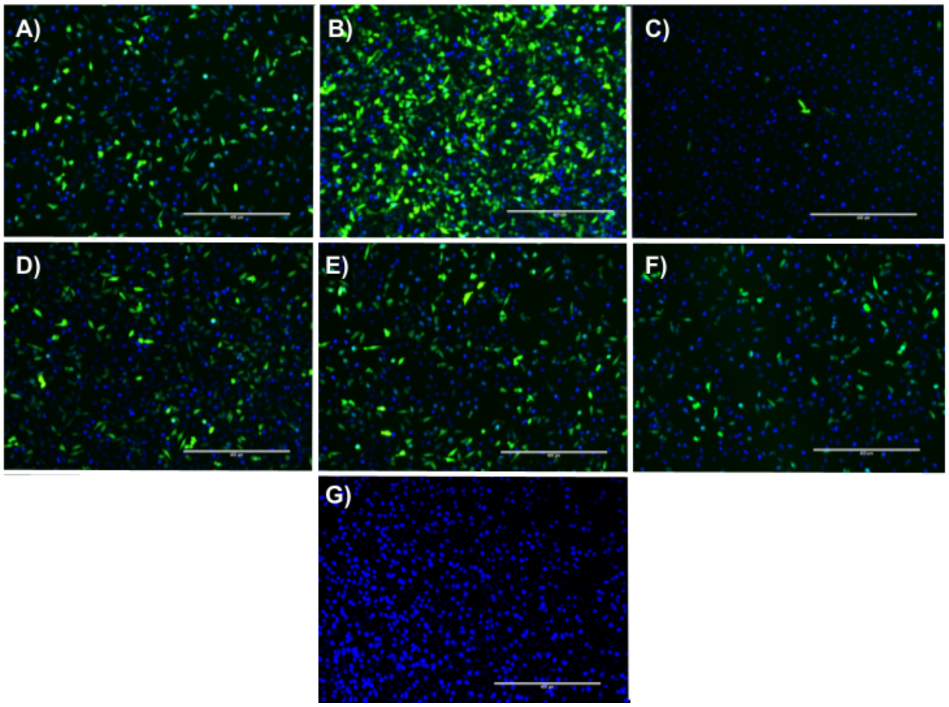


**Supplementary Figure 14. Sample of HeLa Energy Density EQ**.

Through 100µl electroporations using Opti-MEM with 4mm and 2mm cuvettes, the effects of pulse duration when energy density is controlled was examined. Using an energy density constant of 48 KJ/L, various parameters were derived that varied with respect to voltage and duration. Cells were fixed with 4% PFA and stained with 10µM DAPI(Blue). The Experimental conditions were as follows: **A)** 4mm 380 V(950 V/cm) 4ms **B)** 2mm 120 V(600 V/cm) 10ms **C)** 4mm 100 V(250 V/cm) 57.6ms **D)** 4mm 200 V(500 V/cm) 14.4ms **E)** 4mm 240 V(600 V/cm) 10ms **F)** 4mm 280 V(700 V/cm) 7.3ms **G)** Control. Scale = 400µm.

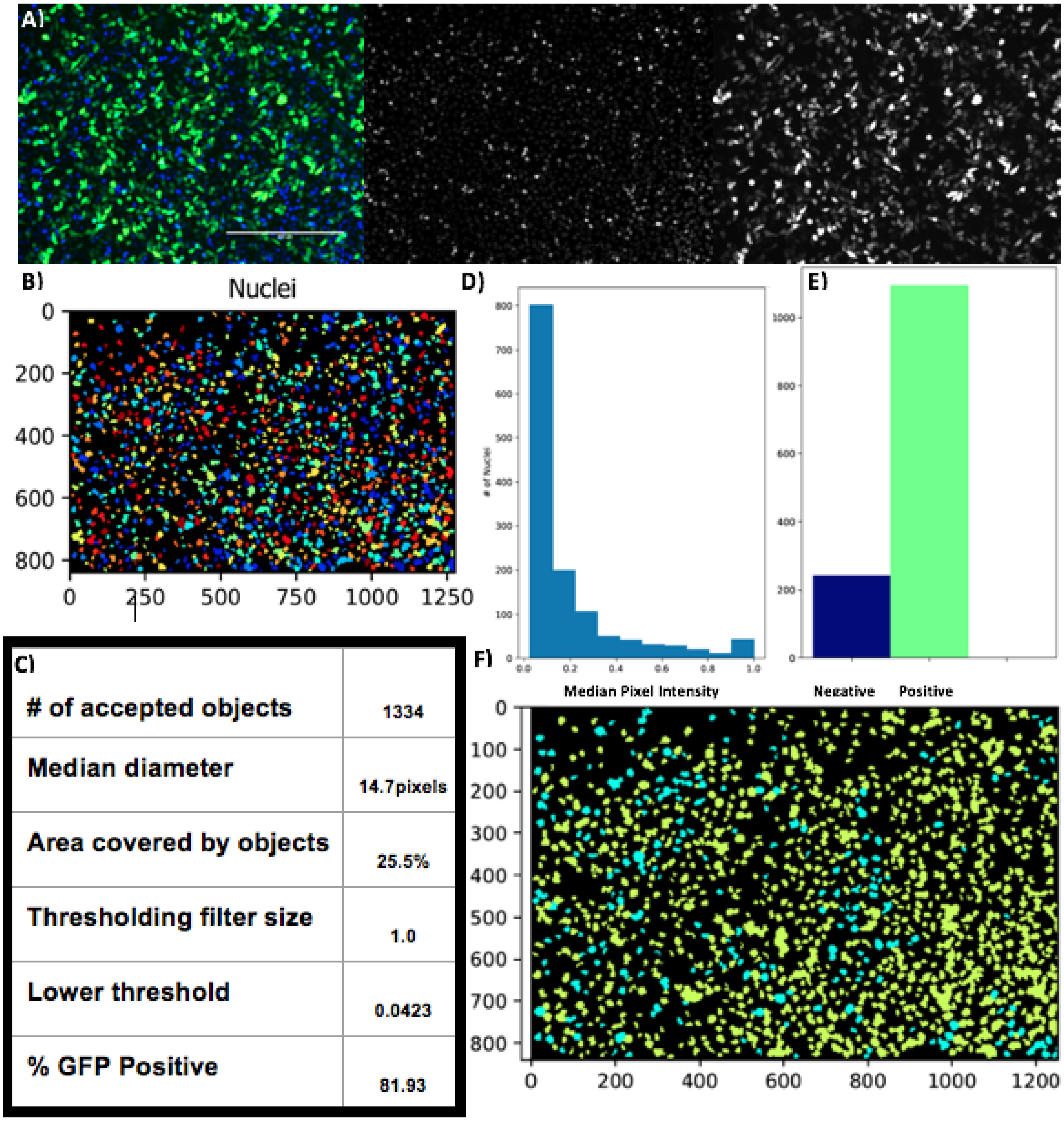


**Supplementary Figure 15: Summary of CellProfiler Analysis Pipeline With HeLa Cells As Example.**

Using a 2mm cuvette, this experimental condition was electroporated at 120V for 10ms. **A)** The RGB images was split into greyscales for the respective channels (GFP and DAPI). **B)** The DAPI channel grayscale image is used to identify the cell's nucleus **C)** and record number and size information to produce an image masking composite. The location of the nuclei are then mapped and overlaid onto the GFP channel grayscale image. **D)** The distribution of GFP channel pixel intensities in the areas occupied by the mapped nuclei are recorded; **E)** and used to classify the cells as GFP positive or negative depending on where they fall with regard to the GFP classification threshold (0.0423 for this image). This minimum threshold is determined empirically, using a control image to identify the minimum pixel intensity required to eliminate false positive nuclei. **F)** The classification information is then overlaid onto the nuclei mask composite to visualize the distribution of positive cells(yellow) and negative cells (light blue).

I. $W = \sigma E^2 t \upsilon$

XII. $\frac{W}{(\sigma)(t)(\upsilon)} = E^2$

XIII. $\sqrt{\frac{W}{\sigma t \upsilon}} = \sqrt{E^2}$

XIV. $\sqrt{\frac{W}{\sigma t \upsilon}} = E$

X. $\Delta\Phi = \frac{3}{2} Ercos\theta$

XI. $\Delta\Phi = \frac{3}{2}\sqrt{\frac{W}{\sigma t \upsilon}} rcos\theta$

**Supplementary Figure 16: The transmembrane potential can be related to the energy density of electroporations.** Energy density (I), and the Schwan equation (X) can be combined to predict a possible relationship between energy density, and cellular transmembrane potential Equations(XII to XIV, and XI). Here, ΔΦ represents transmembrane potential, *r* represents cell radius, θ represents the angle of the cell with respect to the direction of the electric field, *W* represents pulse power, σ represents sample buffer conductance, *t* represents time, and *υ* represents sample buffer volume.
